# A Weights-based variant ranking pipeline for familial complex disorders

**DOI:** 10.1101/2023.08.14.553248

**Authors:** Sneha Ralli, Tariq Vira, Carla Daniela Robles-Espinoza, David J. Adams, Angela R. Brooks-Wilson

## Abstract

Identifying genetic susceptibility factors for complex disorders remains a challenging task. We have developed a weights-based pipeline to prioritize variants and genes in collections of small and large pedigrees where genetic heterogeneity is likely, but biological commonalities are plausible. The **W**eights-based v**A**riant **R**anking in **P**edigrees (WARP) pipeline prioritizes variants using 5 weights: disease incidence rate, number of cases in a family, genome fraction shared amongst cases in a family, allele frequency and variant deleteriousness. Weights, except for the population allele frequency weight, are normalized between 0 to 1. Weights are combined multiplicatively to produce family-specific-variant weights that are then averaged across all families in which the variant is observed to generate a multifamily weight. Sorting multifamily weights in descending order creates a ranked list of variants and genes for further investigation. WARP was validated using familial melanoma sequence data from the European Genome-phenome Archive. The pipeline identified variation in known germline melanoma genes *POT1, MITF* and *BAP1* in 4 out of 13 families (31%). Analysis of the other 9 families identified several interesting genes, some of which might have a role in melanoma. WARP provides an approach to identify disease predisposing genes in studies with small and large pedigrees.

## Introduction

Next-generation sequencing can detect common and rare genetic variants and has been proven key to identifying disease-causing mutations in families affected by Mendelian or complex disorders. Germline variants involved in Mendelian disorders can be detected by searching for variants that segregate in a highly penetrant manner in one or more families [1,2]. Disease gene identification often involves filtering or ranking variants using information such as functional impact, predicted pathogenicity, variant conservation status, and/or allele frequency. Filtering approaches that use hard cut-offs may discard disease-causing variants; a problem for complex disorders and those with incomplete penetrance, where variant functional effect may be less impactful and disease alleles not as rare as for Mendelian disorders. Synonymous variants, non-coding variants or variants with no allele frequency recorded in the public databases may be excluded during some filtering processes. Synonymous changes have been implicated in human diseases, however, by affecting splicing and mRNA stability and altering protein conformation [3]. Similarly, non-coding variants have been found to increase the risk of some diseases [4,5]. Ranking, rather than filtering, places variants on a continuum and allows subsequent choice of subsets of variants for further examination and replication.

Various tools, approaches and pipelines have been developed to either rank or filter variants to aid in detecting putative disease-causing genes that functional studies can later verify. VariantDB filters on parent-offspring and sibling relationships to enable filtering for mode of inheritance such as *de novo*, dominant or recessive. It then filters on variant information such as population-based variant allele frequency, pathogenicity and function [6]. It relies on Mendelian inheritance models and pedigree data limited to parent-offspring and sibling relationships. KGGSeq is a tool that can be used for both Mendelian and complex disorders by filtering of variants on a disease inheritance model, shared identity-by-descent segments or allele frequency followed by pathogenic variant prediction [7]. Following the filtering step, KEGGSeq performs biological analysis of filtered variants at gene, pathway, protein interactions and phenotype level using a novel bit-block encoding algorithm that aids in faster analysis. [7]. The filtering of variants using genetic inheritance model or shared segments is skipped and variants are directly analyzed for functional score when analysing complex disorders[7]. Both tools ignore information such as disease age of onset or family-based details such as extent of genome shared between affected cases in a family.

Some approaches use variant segregation to rank or filter variants within a pedigree. A pipeline called the Familial Cancer Variant Prioritization Pipeline (FCVPP) version 1 identifies germline variants based on variant segregation and prioritizes them using CADD scores that are later evaluated on the conservational score, damage prediction, and the predicted functional effects of variants [8]. An upgraded version of the FCVPP (version 2) also prioritizes regulatory germline variants [9]. FCVPP performs best with a large pedigree with sequenced affected and unaffected family members, though FCVPP version 2 can be applied to trio pedigrees. Neither version of FCVPP can be applied to a group of small and large families. MendelScan assigns scores based on segregation, allele frequency, variant functional effect, and gene expression to rank variants that can be narrowed down to identify disease-causing haplotypes; it works well on autosomal dominant disorders [10]. Reliance on variant segregation can be hampered by incomplete penetrance and genetic heterogeneity in complex disorders.

Tools have also been developed that work best with small pedigrees. Var-MD analyzes a set of exome variants by first filtering on the Mendelian mode of inheritance and then generates a ranked list of potential disease-causing candidates based on pathogenicity, population frequency, genotype call quality, and sequence coverage [11]. Another tool, pVAAST (pedigree-Variant Annotation, Analysis and Search Tool), uses a statistical framework that integrates linkage analysis, association analysis and functional variant prediction [12]. This tool overcomes incomplete penetrance and locus heterogeneity for linkage analysis but works best with small families with rare Mendelian diseases or requires large families for common complex diseases. Requena, Gallego-Martinez and Lopez-Escamez (2017) developed an approach that can be applied to small pedigrees, which combines multiple tools such as the PAVAR score, Variant Annotation Analysis and Search Tool (VAAST-Phevor), Exomiser-v2, CADD, and FATHMM to identify candidate variants [13]. This approach is limited to autosomal dominant disorders and combining different tools for variant lists might lead to the loss of putative disease-causing variants. These tools/approaches, therefore, cannot be applied to a mix of large and small families for a complex disorder.

Most of these tools/pipelines/approaches focus on whole exome sequence data, although some incorporate features that can evaluate non-coding and regulatory variants. The drawback of these tools is that some rely on the mode of inheritance, work well with either small or large pedigrees but not both, and often ignore family-based information such as genetic sharing amongst the cases in a family.

We have developed a **W**eights-based v**A**riant **R**anking in **P**edigrees (WARP) pipeline to overcome the limitations of the existing tools and approaches, particularly for analysis of collections of small and large pedigrees with complex genetic disorders, where genetic heterogeneity is likely but biological commonality is plausible. Our pipeline ranks variants by applying five weights. The five weights are based on (a) age of diagnosis or rarity of disease in the cases, (b) the total number of cases in a family, (c) genome fraction shared amongst sequenced cases in a family, (d) population allele-frequency and (e) variant deleteriousness. These weights are combined for each family to generate a Family Specific variant weight (FSVW). Obtaining a ranked list of variants for a group of families is accomplished by generating a multifamily weight (MFW) by taking an average of the FSVW of the families in which the variant is observed. The MFW are ranked in descending order and then analyzed for biological commonalities.

This pipeline has several advantages over the existing tools and approaches. It ranks variants using family and variant-based information. Age of diagnosis is integrated, which gives greater weight to earlier onset cases that are more likely to have a genetic basis, as opposed to environmental or lifestyle-based cases that often develop later. Cases from large and small families can be analyzed jointly, maximizing the amount of data that can be combined for understanding the disease etiology. The pipeline can incorporate data from distant family members such as second-degree and third-degree relatives, which reduces the number of shared variants and decreases the search space for a given family. The modular design of the pipeline also provides effortless updates of component databases such as CADD.

Here we demonstrate this pipeline on exome data from melanoma families obtained from European Genome-phenome Archive (EGA) EGAS00001000017. Robles-Espinoza et al. (2014) studied the families from this dataset, and identified two *POT1* variants in two families [14].

## Materials and Methods

### Families

The melanoma families are part of the sequence data deposited at the European Genome-phenome Archive (EGA), which is hosted by the EBI and the CRG, under accession number EGAS00001000017, and have been previously published [14]. The dataset includes exome data from 89 melanoma cases. Of the 89 cases, 32 belong to melanoma families where more than one case was sequenced; they are part of 13 melanoma families that were used for the analysis by WARP. The rest belong to melanoma families with only one case sequenced (43 cases) or single cases that presented with multiple primary melanomas, multiple cancers, or an early age of onset (14 cases).

### Sequence Alignment, Variant Calling and Quality control

The EGAS00001000017 dataset provides downloadable bam or srf files for the 89 melanoma cases. For cases where .srf files are available, they were first converted into fastq files using srf2fastq [15]. The fastq files were mapped against the human reference genome (GrCh37) utilizing Burrows-Wheeler Aligner mem version 0.7.6a [16]. Aligned reads were filtered and sorted using sambamba version 0.5.5 [17], and the BAM files generated used in the variant discovery process performed using GATK’s (version 4.0.2.1) HaplotypeCaller (32). Variants were then jointly called to generate a single VCF file.

The VCF file containing single nucleotide variants (SNV) and insertions and deletions (indels) was filtered for variant quality using Variant Quality Score Recalibration (VQSR) with truth sensitivity at 99.0% for both SNVs and indels. Variants that did not pass VQSR were removed. Multiallelic sites were converted into biallelic sites with left alignment and normalization of variants was performed using Bcftools [18]. The VCF file containing all the cases was split into individual family VCF files, and positions with missing genotypes were removed. As VQSR uses a site-specific approach, the quality filter of GQ≥20 & DP≥8 was applied.

The WARP pipeline includes a step that removes variants found in a sequenced informative unaffected individual (for example, the unaffected parent in a family where the disease is clearly not transmitted through that parent). For each family, the shared genotype of 0/1 or 1/1 is retained in the VCF file. In the case of an unaffected sequenced individual, the genotype 0/1 or 1/1 positions shared with the unaffected individual would be removed. For the melanoma families used here, only case sequences were available, and so this step was skipped.

### Annotation

For each family, the shared alternate allele set was annotated using Snpsift version 5.0 [19] and VEP version 103 [20]. The non-Finnish European allele frequency field was annotated from Genome Aggregation Database (gnomAD 2.2.1) using Snpsift. ExAC and 1000Genomes European allele frequency were annotated using VEP. Combined Annotation Dependent Depletion (CADD) RawScores and Phred Scores were annotated using CADD version 1.6 [21] from the website (https://cadd.gs.washington.edu/score). All fields from dbSNP version 155 (21) were used for annotation using Snpsift.

### The Weights based pipeline

Annotated shared variants from multiple families were analyzed by weighting them on five criteria. These criteria are individual weight (IW), family weight (FW), sharing weight (SW), population allele frequency weight (PAFW), and prediction weight (PW).

IW for sequenced affected individuals in a family for this study is derived from an open-access tool CancerData (https://www.cancerdata.nhs.uk/incidence_and_mortality), published by the National Cancer Registration and Analysis Service. Data extracted from CancerData includes melanoma cases for a 5-year period from 2015-2019 in combination with age and sex. The incidence rates are reported per 100,000 age-standardized rates. The file with incidence rates for all age groups and sex for melanoma is referred to as the master file. For each sequenced affected individual in a family, IW was computed by extracting the incidence rate using their age and gender from the master file. An inverse incidence rate is taken to upweight variants in cases diagnosed with melanoma at a younger age. IW is normalized to the range of 0-1 by taking the ratio of the IW incidence rate of an individual to the maximum incidence rate observed in the familial dataset. As two or more affected individuals are analyzed in each family, an average of the normalized incidence rate for affected individuals in a family is assigned to all the shared variants of a family.

FW is based on the number of affected individuals in a family regardless of their sequencing status. For instance, if a family has five affected individuals, of which two are sequenced, then the FW assigned for this family is five. The rationale for FW is based on the fact that families with more affected individuals are more likely to have a genetic basis. In contrast, in some small families, the disease may be caused by the coincidental occurrence of sporadic disease rather than the segregation of a susceptibility gene(s). Variants shared by a family are given the same FW; thus, a higher weight is given to families with a greater number of affected individuals. FWs are normalized to the range of 0-1. Normalization is performed by dividing the number of cases in a family by the maximum number of affected individuals observed in a single family in the dataset.

SW is the inverse of the fraction of genomic sharing between the sequenced individuals of a family. Families with higher numbers of informative DNA samples allow us to rule out more variants observed in family members by requiring that they are shared between affected relatives to be of interest. Shared variants in a family have the same SW. This weight is normalized to the range of 0-1. Normalization is performed by taking the ratio of SW in a family to the maximum SW observed in the dataset.

PAFW uses allele frequency in population cohorts, such as ALFA [22], gnomAD [23], ExAC [24], and 1000Genomes [25], which indicates the rareness of a variant. Both rare and common variants are considered rather than applying an arbitrary cutoff to filter variants out. This weight is calculated as (1–allele frequency), giving higher weight to rare variants. When a variant frequency is identified in only one database, then that database is used to obtain its PAFW. In contrast, if a variant is observed in more than one database, then the PAFW is calculated from one of the databases in a preferred order based on the sample size of the database. Preference is given to allele frequency from ALFA, gnomAD, ExAC, and then 1000Genomes as ALFA database has the largest sample size followed by gnomAD. Variant frequencies not identified in any of the databases are assigned an arbitrary low allele frequency, assuming that these variants are rare and not yet discovered in existing allele frequency databases.

PW is applied to variants based on their predicted deleteriousness. The rationale for this is its capability to up-weight damaging variants. CADD raw scores are used to calculate the PW as it provides a range of relative differences in deleteriousness amongst the variants [26]. Raw scores are obtained from the CADD database, where a higher raw CADD score indicates that a variant is more likely to have deleterious effects. Negative CADD raw scores are converted into low-value positive scores so they can be combined with other weights. All negative values are given a value of 0.000001, which is lower than the smallest positive value in the familial dataset. PW is normalized to 0-1 by taking the ratio of PW for a variant to the maximum PW in familial dataset.

Each variant received these 5 individual weights, each between 0 and 1. These 5 weights were combined multiplicatively to generate a family-specific variant weight (FSVW) for each variant in a family outputted into a .tsv file. These files with FSVW are converted into .tab files and annotated to individual family VCF files using the BCFtools annotate command [18]. Once each VCF file is annotated with FSVW, they are merged using BCFtools [18], which generates a single VCF file. Variant details from the merged file, such as allele frequency, CADD score, gene name, and FSVW were extracted using the BCFtools +split-vep plug-in [18] to generate multifamily weight (MFW). For each variant in the extracted file, FSVWs were averaged to generate an MFW. The average is based on the number of families in which the variant is observed, as these families are expected to show genetic heterogeneity, so some families might have the variant while others may not. Therefore, MFW is generated by taking the average instead of giving higher weight to variants observed in multiple families. The number of families harboring the variant is also calculated, which is used while interpreting the variant of interest. The multifamily weight is sorted in descending order generating a ranked variant list. Code for the WARP pipeline can be found at https://github.com/s-ralli/WARP.git

### Analysis of variants for biological commonalities

Assessment for biological commonalities was done by examination of top-ranked variants from each family, and through literature searches. Common variants (allele frequency > 0.01) that were highly ranked in the melanoma families data set were examined in the GWAS catalogue [27]. Variants of interest were visually inspected using Integrative Genomics viewer(IGV) version 2.4 [28] and excluded if deemed artifacts.

Starting at the top of each family’s ranked list of variants, rare variants (allele frequency ≤ 0.01) were checked and those that were either in pseudogenes or that did not withstand a TraP cut-off score of 90 percentile [29] for synonymous variants, or that were deemed artifacts upon examination in the Integrative Genomics Viewer (IGV), were excluded. The process was repeated with each family’s ranked list of variants until 15 verified, highly ranked variants were identified. A gene set generated from the 15 most highly ranked rare variants from each family was analyzed using the gene set/Mutation analysis tool of the Reactome Functional Interaction (FI) in Cytoscape version 3.9.1. For this purpose, the 2021 ‘ReactomeFI Network’ dataset option was used to create interaction networks without adding any linker gene. Enrichment analysis for the networks was performed using the Analyze network functions for the pathway or GO Biological processes. The p-values were calculated based on binomial test and the adjusted p-value ≤0.05 was considered significant. The adjusted p-value is computed by ReactomeFI using the using the Benjamini-Hochberg method.

Literature-based biological commonalities analysis was performed by looking for rare and common variants in genes previously known to have germline mutations in melanoma families, genes known to be somatically mutated in melanoma tumors, and genes identified in GWAS studies of melanoma. The source for known germline melanoma genes was Toussi, A et al. (2020) [30]; somatically mutated genes were acquired from the COSMIC cancer gene census (https://cancer.sanger.ac.uk/census) [31], and GWAS genes were taken from the GWAS catalogue [27].

## Results

The WARP pipeline is summarized in Fig 1. We anticipate that a disease-causing mutation would be shared amongst the cases of a family. The pipeline was validated on exome sequence data from 13 families with 32 cases in the EGAS00001000017 dataset. These families were verified by KING version 2.1.8 [32] relationship inference. There were, in total, 91,021 variants, of which 86,639 (95.2%) are common variants (allele frequency >0.01), 4,107 (4.5%) are rare variants (allele frequency ≤0.01), and 279 (0.3%) are novel variants with no allele frequency reported in public datasets.

**Fig 1:**
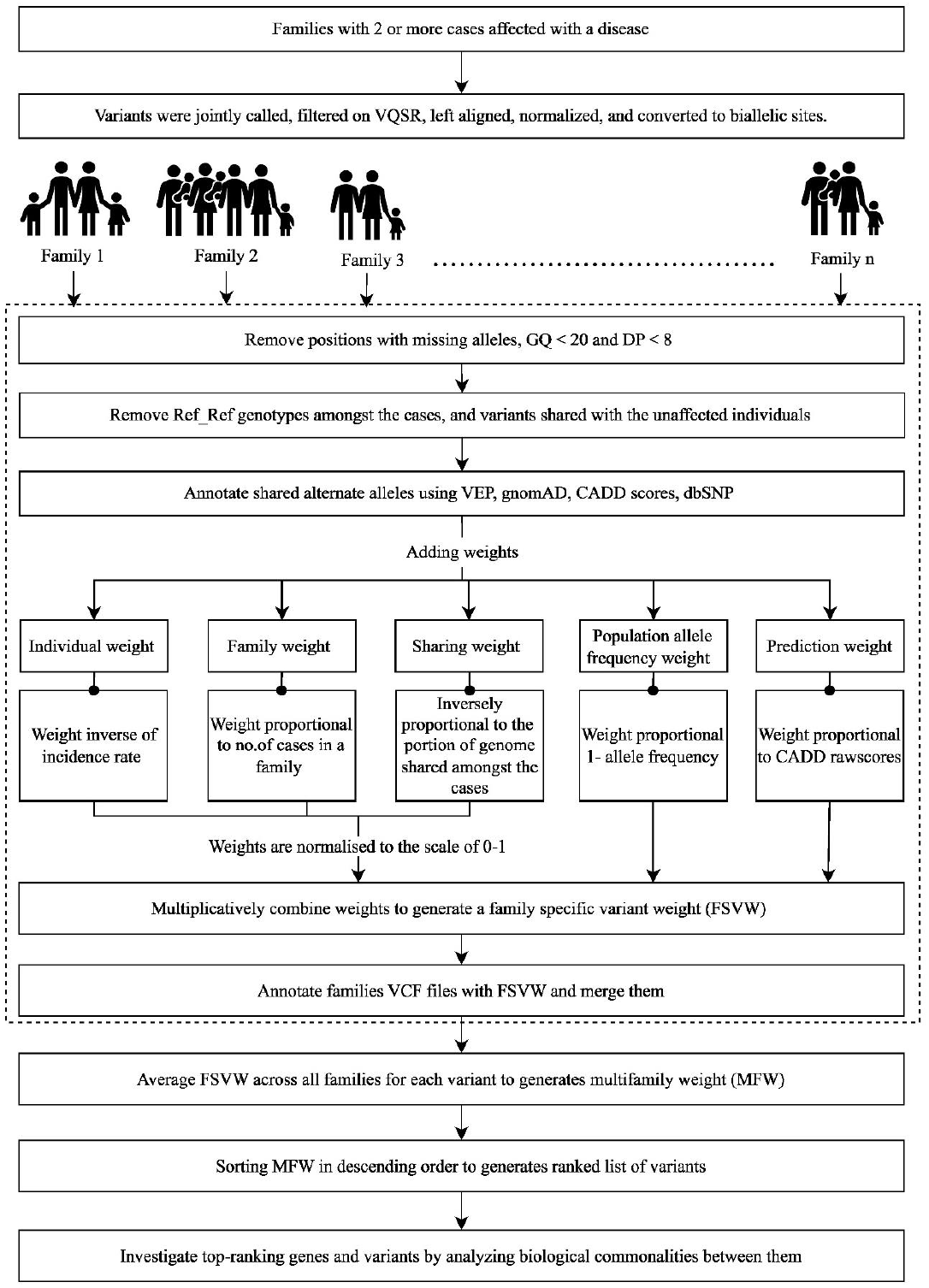
Overview of the weight-based variant ranking pipeline for complex familial disorders. Generated using draw.io (version 21.6.5; https://app.diagrams.net/)

### Rank of previously identified POT1 variants

A published paper reported two *POT1* variants rs587777472 (g.124503684T>C) and rs587777473 (g.124465412C>T) that are observed in two families. The rs587777472 is a missense variant that alters the amino acid from tryptophan to cysteine in the highly conserved N-terminal oligonucleotide-/oligosaccharide-binding (OB) domain of the POT1 protein observed in 5-case family UF20. In the combined ranked list for all families together, this variant is at position 37 (99.96 percentile). The second variant, rs587777473, is a stop gain variant observed in 6-case family AF1. This variant is ranked at the 96.75 percentile (ranked 2955th). Both variants are predicted to be damaging or probably damaging by SIFT [33] and Polyphen [34].

### Assessment of biological commonalities among top-ranked genes

The top 15 rare variants were chosen from each of the 13 families, which resulted in a set of 194 variants in 188 genes. A deletion variant, rs199851144 in *FAM111B*, was observed in two families, UF19 and UF21. Pairs of different variants in the same genes were identified in 5 genes (Table 1). Two variants, rs117307819 and rs17304212 were observed in two or more families, but each one made it to the top 15 in only one family. rs117307819 in *ELAVL1* is a synonymous variant with a CADD score of 12.75. The TraP score for this variant is 0.287, which is above the top 92.5th percentile in TraP, indicating that it is predicted to be possibly damaging. The rs117307819 variant is observed in three families – UF16, UF20, and UF10. Variant rs17304212 is a missense variant in *DFNB59* with a 23.9 CADD score and conflicting interpretations of pathogenicity on Clinvar. This variant is observed in two families, NF2 and UF1.

**Table 1:** Summary of 5 genes with more than one variant identified during the biological commonalities of top 15 analysis in 13 families.

| Gene | ID | Allele Frequency | CADD score | Consequence | Family IDs | Rank (percentile) |
| --- | --- | --- | --- | --- | --- | --- |
| *DNAH11* | rs775108833 | 3.76E-05 | 24.7 | Missense variant | UF19 | 99.91 |
|  | rs72657389 | 3.29E-03 | 26 | Missense variant | UF7 | 99.81 |
| *EFHB* | novel | novel | 28.7 | Missense variant | UF21 | 99.79 |
|  | rs145933876 | 6.64E-03 | 23.9 | Missense variant | UF10 | 99.76 |
| *POT1* | rs587777473 | 9.13E-06 | 33 | Splice acceptor variant | AF1 | 93.29 |
|  | rs587777472 | 5.96E-05 | 24.4 | Missense variant | UF20 | 96.75 |
| *RYR3* | rs201633381 | 1.01E-03 | 25.6 | Missense variant | UF19 | 99.96 |
|  | rs190035689 | 1.42E-04 | 25 | Missense variant | UF19 | 98.38 |
|  | rs181264765 | 2.73E-03 | 32 | Missense variant | NF3 | 99.75 |
| *USH2A* | rs80338902 | 1.55E-03 | 28.9 | Missense variant | UF15 | 99.13 |
|  | rs1356404884 | 1.76E-05 | 23.1 | Missense variant | NF3 | 98.38 |

The top 15 variant set also included both the variants in *POT1* found in family UF20 and AF1. In addition to the *POT1* variants, two other families have variants in known germline melanoma genes, *BAP1* and *MITF*. Family NF3, had a novel frameshift variant g.52436841T>TAA (CADD score of 33) in *BAP1*. This variant is shared by all four sequenced cases and is ranked at position 121 (99.87 percentile) by the pipeline. The *BAP1* variant was not reported by Robles-Espinoza et al. (2014); however, another study performed on the same family using new generation aligners and callers did identify the *BAP1* variant [14,35]. UF10 is a small family with 3 sequenced cases showing a missense variant rs149617956 that changes the amino acid from glutamic acid to lysine in known melanoma gene *MITF*. This variant is reported to be pathogenic/likely pathogenic in Clinvar and is ranked at position 782 (99.14 percentile) by the pipeline.

Given that four families had variants in known germline melanoma predisposing genes, we re-ran the pipeline excluding these four families, and including just the nine families with no variant in a known germline melanoma gene. The top 15 rare variants from each family were then selected for these nine families based on the MFW; the set included 135 variants. The identified variants and genes were investigated through literature search to get insight into their role in melanoma. Potential melanoma genes for these 9 families after the review are summarized in Table 2.

**Table 2:**
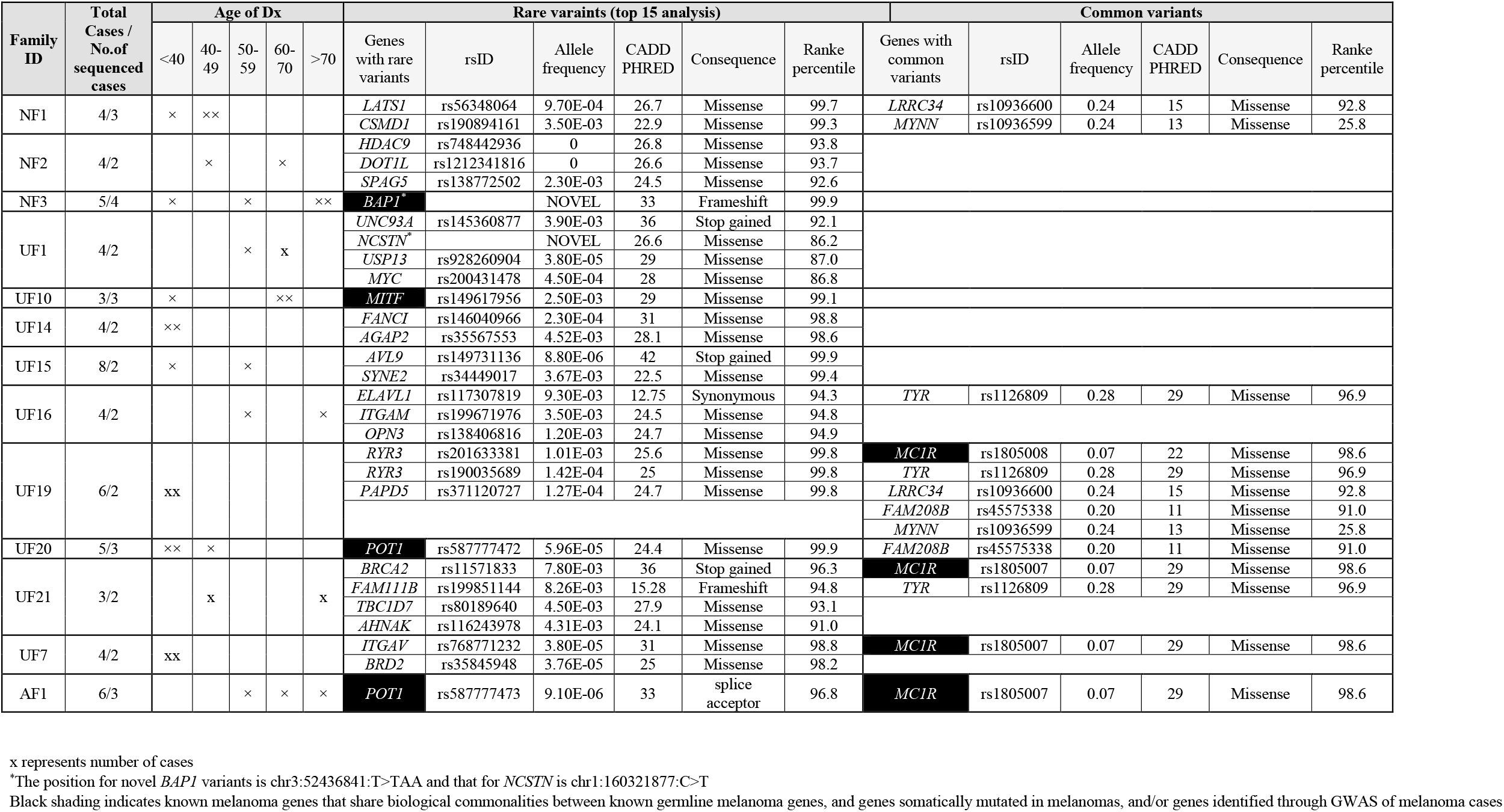
Putative melanoma susceptibility genes with rare and common variants detected in 13 melanoma families.

188 genes are represented in the 194 variants in the top 15 variant set of 13 melanoma families and querying them on ReactomeFI led to the generation of 12 networks with 45 genes (Fig 2). 22 pathways are enriched with adjusted p-values ≤0.05 in these 45 genes. Table 3 summarizes the enriched 3 pathways from Reactome, 3 from KEGG, 10 from NCI PID, 5 from Biocarta and one from Panther. The top two pathways in the network-based analysis are Regulation of retinoblastoma protein and Beta2 integrin cell surface interactions from the NCI PID database with an adjusted p-value of 0.0146. The GO biological processes are enriched by 309 processes within these 45 genes where the FDR value is ≤ 0.05. These 309 processes include sets of 8 GO biological processes where the number of genes in the process is > 200 and 241 GO biological processes were the number of query hit genes is 1. Table 4 summarizes the top 10 GO biological processes identified by the network based ReactomeFI. The top GO biological process is melanocyte differentiation, with an FDR of 0.02.

**Table 3:**
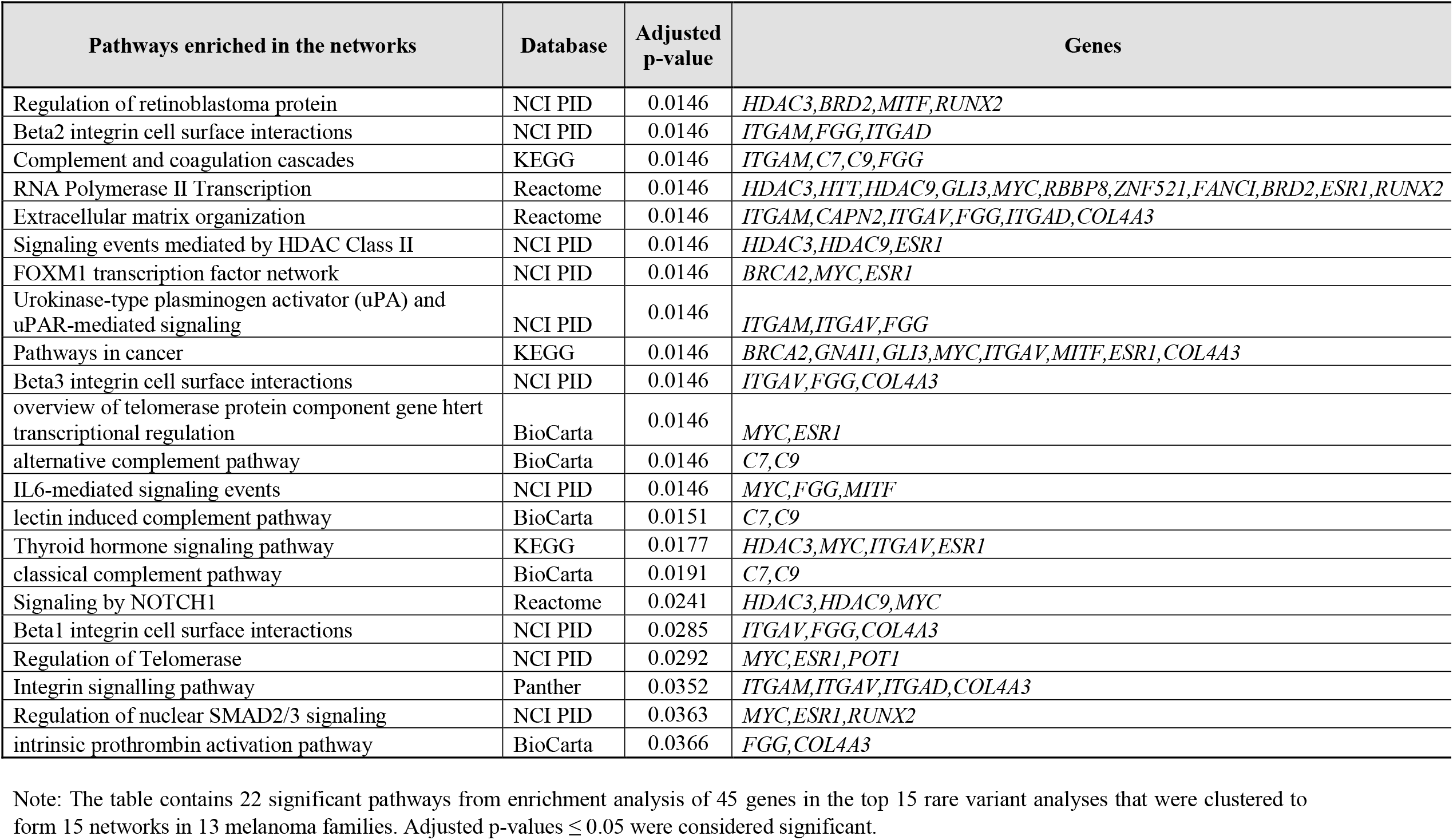
Pathway enrichment analysis with ReactomeFI.

**Table 4:** Top 10 GO Biological Processes identified through enrichment analysis

| GO Biological Processes | Adjusted p-value | Genes |
| --- | --- | --- |
| melanocyte differentiation | 0.0202 | *USP13, GLI3, MITF* |
| positive regulation of protein localization to cell cortex | 0.0241 | *GPSM2, GNAI1* |
| positive regulation of spindle assembly | 0.0241 | *GPSM2, SPAG5* |
| establishment of protein localization to telomere | 0.0261 | *BRCA2, POT1* |
| cell adhesion mediated by integrin | 0.0284 | *ITGAM, ITGAV, ITGAD* |
| cell-matrix adhesion | 0.0284 | *ITGAM, ITGAV, FGG, ITGAD* |
| protein import into peroxisome matrix | 0.0361 | *PEX2, PEX6* |
| cell division | 0.0361 | *GPSM2, GNAI1, RBBP8, KNTC1, SPAG5, LATS1* |
| G1/S transition of mitotic cell cycle | 0.0361 | *MYC, RBBP8, LATS1* |
| cellular response to estrogen stimulus | 0.0361 | *MYC, ESR1* |

Analyses included 15 ReactomeFI networks identified from the top 15 rare variant analysis set from 13 melanoma families.

**Fig 2:**
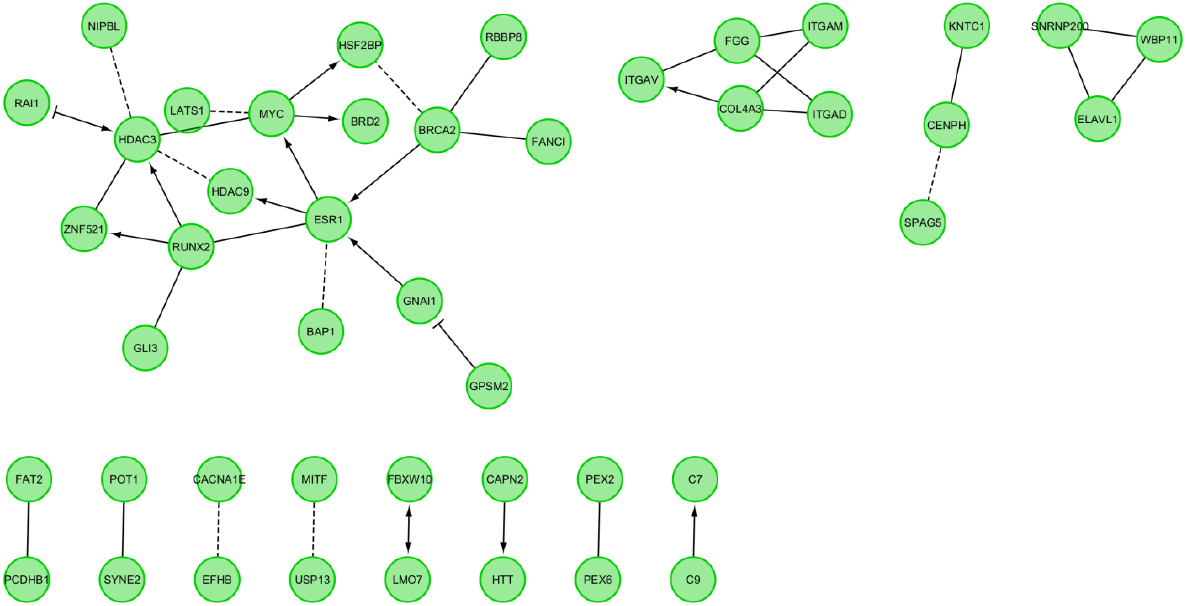
Biological interaction network generated using Cytoscape v3.9.1 for top15 variants from 13 melanoma families.

### Literature-informed assessment of biological commonalities

Genes known to have germline mutation in melanoma cases, genes known to be somatically mutated in melanoma tumors, and genes identified through melanoma GWAS, and their overlaps, are summarized in Fig 3. Three genes, *MITF, CDKN2A*, and *TERT*, are shared between all three categories. Only one gene, *MITF*, had a rare variant rs149617956 in family UF10. *CDK4* and *BAP1* are in both the somatic and germline categories. No variants in *CDK4* were present in the melanoma families, but one rare variant, g.52436841T>TAA in *BAP1*, was identified in family NF3. One gene, *MC1R*, is shared between the GWAS and germline categories. Two common variants rs1805007 (g.89986117C>T) and rs1805008 (g.89986144C>T) in *MC1R* is associated with melanoma in the GWAS catalogue. The *MC1R* variant rs1805007 is associated with freckling and sun sensitivity and was present in the AF1 family previously shown to have a *POT1* variant [14] and had an odds ratio (95% confidence interval) of 4.38 (2.03–9.43) for the effect allele T [36]. This variant is also found in two additional families, UF21 and UF7, which did not have a variant in known melanoma genes. One of the cases from UF7 is homozygous for the T allele. Another *MC1R* variant, rs1805008, was observed in family UF19. The effect allele/non-effect allele for the rs1805008 is T/C, and the family is heterozygous for the T effect allele. This allele has an odds ratio (95% confidence interval) of 1.64 (0.85–3.19) with an effect allele frequency of 0.098 [36].

**Fig 3:**
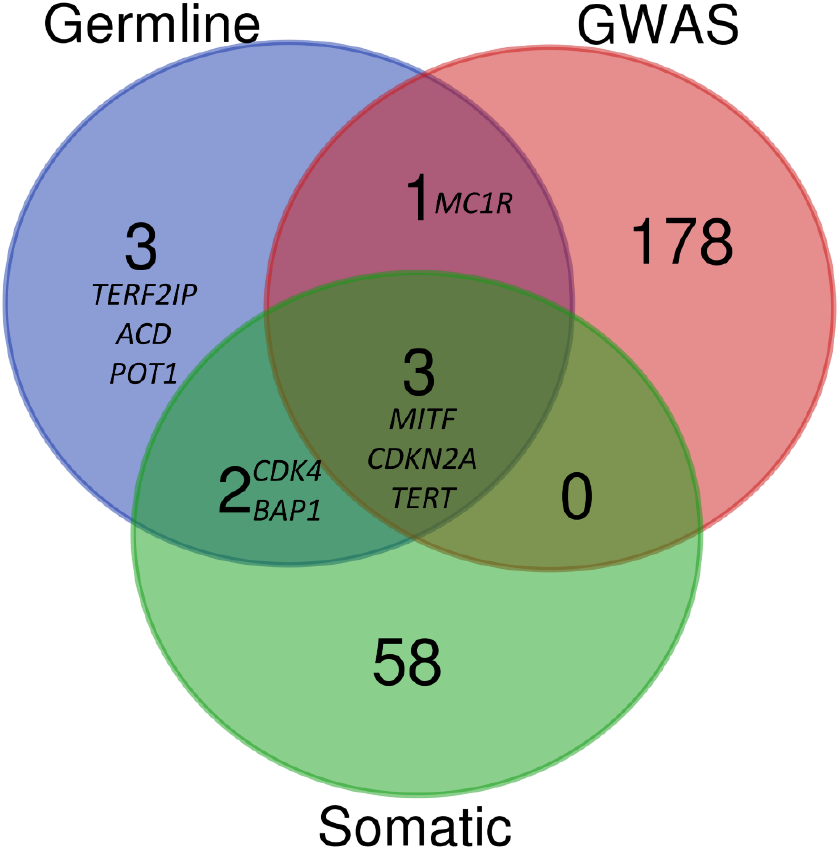
Biological commonalities between known germline melanoma genes, and genes somatically mutated in melanomas, and/or genes identified through GWAS of melanoma cases. Generated using Venn Diagrams (https://www.vandepeerlab.org/?q=tools/venn-diagrams)

The entire ranked list of variants was searched for rsIDs associated with melanoma in the GWAS catalogue. Known GWAS variants were identified in the seven genes *DSTYK, TYR, FAM208B, LRRC34, MYNN and MC1R*. Table 2 lists the common variants, their percentile rank, and the family they are present in. The effect allele/non-effect allele for rs3851294 in *DSTYK* is A/G, and the effect allele frequency is 0.098. The germline exome data from the melanoma cases show that the cases in family UF1 are heterozygous for the effect allele A. In a previous study, this variant had an odds ratio (95% confidence interval) of 1.05 (1.03-1.07) [37].

The *TYR* variant rs1126809 has effect allele/non-effect allele A/G, and the effect allele frequency is 0.27632 with the odds ratio (95% confidence interval) of 1.27 (1.16, 1.40) [38]. Families UF16, UF19 and UF21 are heterozygous for effect allele A, whereas family UF1 is homozygous for the effect allele. *FAM208B* variant rs45575338, which has been implicated in increased melanoma risk [39], is observed in 3 families – UF1, UF19, and UF20. *LRRC34* variant rs10936600 is present in two families, UF19 and NF1. The effect allele/non-effect allele for *LRRC34* is A/T, and the effect allele frequency is 0.76. The effect allele in these two families is present in the heterozygous effect allele, where the odds ratio for the effect allele is previously reported to be 1.076 [40]. The rs10936599 variant in *MYNN* has been previously identified for melanoma risk variants with the effect allele/non-effect allele is C/T and the effect allele frequency is 0.75 with an odds ratio (95% confidence interval) of 1.06 (1.04-1.08) [37]. This variant is observed in families NF1 and UF19, where the effect allele is heterozygous. Another family, UF14, had one heterozygous case for the effect allele, whereas the other sequenced case was homozygous for the non-effect allele.

## Discussion

A weight-based variant ranking pipeline was developed and validated that aids in the search for variants and genes that affect risk of complex familial disorders. The weights-based pipeline can be used to analyze sets of large and small families together. It works by ranking the variants on the age of diagnosis or rarity of disease subtype of the cases, the number of cases in a family, the genome fraction shared amongst sequenced cases in a family, allele frequency and variant deleteriousness. Ranked variants from large and small families are analyzed for biological commonalities between families and with the known disease literature to identify genes and pathways that may play a role in the genetic etiology of a complex genetic disease.

The pipeline was validated using 13 families from the EGA melanoma dataset EGAS00001000017. No unaffected individuals were sequenced in these families, so it was not possible to reduce the number of variants to be prioritized by removing variants present in unaffected individuals; normally this step would be part of the pipeline.

*POT1* variants rs587777472 and rs587777473 and a common variant rs1805007 in *MC1R* published previously [14] were re-discovered using the pipeline. The pipeline prioritizes the variant rs587777472, which is present in the highly conserved OB domain of the *POT1* protein over the other variant. The variation in the OB domain results in longer telomeres, which predisposes individuals with the variant to develop cutaneous melanoma [14,41]. The rs587777473 variant was of low quality when observed on IGV; however, this variant was validated by capillary sequences in the cases of family AF1 in the previously published paper [14]. *POT1* is a part of the shelterin complex that plays a role in chromosome end maintenance by regulating the length of telomeres [42]. Variants in *POT1* have been detected in various familial melanoma studies [41,43,44], making it a compelling susceptibility gene for familial melanoma. Furthermore, we identified variants in *MITF* and *BAP1*, known germline melanoma genes through our biological commonalities analysis. *MITF* is a transcription factor that plays an essential role in melanocyte differentiation, proliferation and survival by affecting expression of genes such as *BCL2* [45]. rs149617956 variant in *MITF* has been previously identified as a risk variant by linkage analysis of 31 melanoma families under the dominant model with the odds ratio of 2.7 and is involved in increasing the transcriptional activity of *MITF* function by preventing SUMOylating [46]. *BAP1* encodes a tumor suppressor protein that deubiquitinates BARD1 and regulates the E3 ligase activity of the BRCA1–BARD1 complex [47]. *BAP1* has been implicated in uveal melanoma, but studies indicate that this gene can predispose to cutaneous melanoma [48,49]. Interestingly, the rank of variants in known melanoma susceptibility genes indicates that there were many variations with high CADD scores in the genomes of these families. For instance, the top-ranking variant rs149731136 in *ALV9*, plays a role in progression of cell cycle progression [50]. This gene is known to be involved in colorectal cancer and there is a risk of colorectal cancer in families affected with melanoma [51,52] making *ALV9* a candidate susceptibility gene for melanoma. Further, some of these variants in the known melanoma-causing genes might have moved higher in the ranked list after removing false positive variants from IGV.

The additional analysis of 9 families without variation in known germline melanoma genes identified other putative genes that might play a role in melanoma. Family NF1 had a variant rs56348064 in *the LATS1* gene, part of the hippo signaling pathway that acts as a negative regulator of *YAP1*, where inactivation of *LATS1* results in the accumulation of YAP protein and subsequent activation of target cell proliferation genes [53]. Family UF1 contains a variant rs145360877 in *UNC93A*, which was detected in another melanoma family identified through literature search, although little is known about the gene [54]. This family also has a variation rs200431478 in the *MYC* proto-oncogene which is a transcription activator for my genes involved in cell cycle regulation. Copy number variations in *MYC* have been reported in melanoma cases [55]. Family UF21 has a variant rs11571833 that introduces a premature stop codon in *BRCA2*, a gene known to be involved in DNA repair. The premature stop codon identified in family UF21 has been previously reported in another published melanoma family [56]. Recent studies suggest; however, that *BRCA2* may not contribute to pathogenesis of melanoma [57,58]. Another family, UF14, has a variant rs146040966 in *FANCI*, which is part of the Fanconi anemia complementation group. This gene is involved in DNA repair pathway which is known to upregulated in melanoma thereby contributing to melanoma pathogenesis [59]. Family NF2 has a variant rs1212341816 in *DOT1L*, a histone methyltransferase that methylates lysine 79 of histone H3 which aids in the regulation of cell cycle [60]. The role of *DOT1L* has been elucidated in nuclear excision repair (NER) where it recruits NER factors to the site of ultraviolet induced DNA damage [61]. This family and the *DOT1L* variant it carries have been previously reported as co-segregating with melanoma [62]. The role of *DOT1L* in cell-cycle regulation and DNA repair, along with previous mutations reported in this gene, makes *DOT1L* a strong candidate for a susceptibility gene for familial melanoma.

Variants responsible for familial disorders would be expected to be rare in human populations as the variants might be subjected to negative selection. Common variants may impart susceptibility to diseases, but the contribution of these variants is usually small. Both rare and common variants may contribute to complex disorders in families affected with such cancers. For this reason, we designed the pipeline not to filter out any variant but instead rank them, providing an opportunity to evaluate any variant in the ranked list whether rare or common.

We verified the *MC1R* variant reported previously [14]. *MC1R* plays a role in skin pigmentation, protects chromosome stability, and is involved in DNA damage response by increased phosphorylation of DNA repair proteins in melanocytes explaining why variants in *MC1R* are associated with increased risk to melanoma [63,64]. Some families with a common variant rs1805008 or rs1805007 in *MC1R* developed melanoma at a young age, such as in family UF7 with rs1805007 had cases that developed melanoma before age 40. Similarly, the common variant rs1805008 observed in family UF19 had two cases sequenced that also developed melanoma before 40. One of the five genes in family UF19 is *MYNN*, and this same variant is found in family NF1 along with a variant with *LRRC34* known to be associated with an increased risk of melanoma. This family also showed early onset melanoma at the age of < 40 and 40-49 years. The risk of early onset of melanoma might be due to the contribution of the common variants in genes known to be associated with melanoma. These findings would have been ignored if common variants had been filtered out. Remarkably, most of these common variants identified in the EGA melanoma dataset were ranked highly, ranging between 98 to 91 percentiles by the pipeline.

The top pathways detected were the Regulation of retinoblastoma protein and Beta2 integrin cell surface interactions pathway from NCI Pathway Interaction Database. The retinoblastoma pathway plays an essential role in cell cycle control, and dysregulation of the cycle is a hallmark of cancer development. Inactivation of the retinoblastoma pathway has been identified in various cancers, including in the pathogenesis of melanoma [65,66]. Genes in the retinoblastoma pathway that were identified in the melanoma families of this study were *HDAC3, BRD2, MITF*, and *RUNX2*, making them putative candidate genes for melanoma susceptibility. Of these genes, *BRD2* is known to be overexpressed in melanoma and the knockdown of *BRD2* in melanoma cell lines has resulted in cell cycle arrest by preventing the progression of cells from G1 to S phase [67]. The role of *RUNX2* has been evaluated in inducing cell growth, migration and invasion by ShRNA-mediated knock down of *RUNX2* in melanoma cell lines. [68] The role of this gene in tumor progression implies that a germline alteration in *RUNX2* might increase the susceptibility risk of melanoma pathogenesis. Integrins play a role in interconnection of cells with other cells and extracellular matrix. Integrins activate and control many signalling pathways that regulate cell proliferation, migration and apoptosis, indicating that they have a potential role in tumour progression and metastasis in melanoma [69]. Some genes, such as *ITGAM* with the integrin pathway, have been associated with an increased risk of melanoma [70]. Therefore, observing the Beta2 integrin cell surface interactions as a high-ranking pathway is not surprising. Notably, one melanoma family UF7 had germline variations in *BRD2*, part of the retinoblastoma pathway and *ITGAV*, an integrin gene where both the sequenced cases developed melanoma before the age of 40 years. The top GO biological process identified was melanocyte differentiation, where unspecialized cells become melanocytes. Since this process was enriched amongst the genes that contain the most highly ranked variants, it suggests the involvement of *USP13, GLI3* and *MITF* in melanoma.

This pipeline has some strengths and represents significant advances over current approaches to complex disorders. First, variants in a mixture of small and large families can be analyzed together. The large families help filter the shared variants so that the focus is on a few exciting variants; small families can provide a bulk of data in which to seek biological commonalities. Second, the pipeline is not limited to a disease mode of inheritance, which is a requirement for some family-based approaches. The pipeline can therefore be applied to families where the mode of inheritance is unclear or to families affected with complex disorders. Third, the variant databases used in the pipeline can be replaced or combined as better databases for variant prioritization are developed. Fourth, the biological commonalities search provides a unique opportunity to identify novel genes and pathways involved in complex disorders, thereby increasing our knowledge about disease etiology, and analyzing known disease genes with rare and common variants.

The pipeline also has several limitations. There are many common variants, and understanding their impact is generally limited to genome-wide association studies. The pipeline relies on the GWAS catalogue for the analysis of common variants. The function and mechanism of pseudogenes in cancer remain unclear; therefore, the top 15 biological commonalities analysis excludes weighted variants in pseudogenes. The weighted variants in pseudogenes can be analyzed as more information on their function becomes available or if the gene gets classified during reference genome update. The top 15 variants from each family were selected to analyze biological commonalities between families. The number 15 was chosen to allow examination of multiple variants from each family while still being a feasible size of data set to analyze; other cutoffs could be used, depending on the level of genetic heterogeneity suspected of the disease under study. It is expected that the rare variants that might be involved in causing the disease will be ranked higher; the top 15 variant analysis helps investigate a handful of ranked variants from all the families.

Deciphering the genetic architecture of complex disorders is a challenge compared to Mendelian disease, which has had a success rate of about 60-80% [71]. This challenge is in part because complex disorders are multifactorial. Studies of complex familial disorders most often focus on variants in genes and pathways from the literature that are known to be involved in disease etiology. However, focusing on biological commonalities between families allows us to ask this biological question in a way that is less dependent on current knowledge, and has the potential to uncover novel genes and pathways involved in the disease.

## Conclusion

We have developed **W**eight-based v**A**riant **R**anking in **P**edigrees (WARP) pipeline for gene identification in families with complex genetic disorders. The pipeline is able to take advantage of data from both large and small families and is useful in situations where genetic heterogeneity is expected, and biological commonalities are plausible. We validated the pipeline using data from melanoma families in EGA. The pipeline not only detected the *POT1* variants previously reported but also prioritized rare and common variants in other known melanoma-causing genes and identified other genes that may have a role in melanoma. This approach could be applied to sets of families with other complex disorders, particularly cancers.

## Acknowledgements

We thank the Canada’s Michael Smith Genome Sciences Centre (GSC) at BC cancer Sequencing and Bioinformatics platforms for the expert generation of high-quality data. We would also like to thank Nelleke Gruis, Leiden University Medical Center, for providing melanoma families accessed by EGA in this study.

## Author contributions

S.R. and A.R.B-W conceived and designed the experiments. T.V. and S.R. created the computational framework. S.R. planned and carried out analysis of the data. Data curation was done by D.J.A and C.D.E-R. All authors have discussed the results, contributed to writing and approved final version of the manuscript.

## Data Availability

The script for the pipeline to generate FSVW used in the study can be found on GitHub at https://github.com/s-ralli/WARP.git. This work used existing familial melanoma sequence data (EGAS00001000017) obtained from the European Genome-phenome Archive.

